# MATERNAL AUTOPHAGY CONTRIBUTES TO GRAIN YIELD IN MAIZE

**DOI:** 10.64898/2025.12.30.697098

**Authors:** Jie Tang, Tamar Avin-Wittenberg, Erik Vollbrecht, Diane C Bassham

**Affiliations:** Department of Genetics, Development and Cell Biology, Iowa State University, Ames, IA 50011, USA; Department of Plant and Environmental Sciences, Alexander Silberman Institute of Life Sciences, The Hebrew University of Jerusalem, Givat Ram, 9190401, Jerusalem, Israel

## Abstract

Maize is an important crop species that is cultivated worldwide, and increasing maize yield is one of the major goals of plant breeding. The subcellular degradation and recycling pathway known as autophagy is involved in many plant developmental processes including vegetative growth, reproductive development, seed development, and senescence. Here, we demonstrate that maize autophagy-defective *atg10* mutants have delayed flowering and reduced kernel size, weight and number, leading to reduced grain yield. Reciprocal crosses between mutant and parental lines indicate that the maternal plant is the major contributor to the kernel phenotype in *atg10* mutants, rather than the seed genotype. We hypothesize that this may be due to detrimental effects on nutrient remobilization from maternal tissue to seeds during kernel development, leading to reduced yield.

## Introduction

Maize is one of the most important crop species and is cultivated globally. It serves as food, and also as a source for animal feed and industrial raw materials (Godfray et al. 2010). A central goal of maize breeding is to improve yield in order to ensure future food security (Blümmel et al. 2013) and to meet this goal, a better understanding of mechanisms controlling plant growth and seed production is essential.

Macroautophagy (commonly referred to as autophagy) is a degradation and recycling pathway for cellular components, and is conserved throughout eukaryotic species. In plants, it functions to remove damaged proteins or organelles and to recycle nutrients under stress conditions or in response to developmental cues, thereby enabling plant survival and growth (Agbemafle et al. 2023, 2025). Upon the activation of autophagy, the degradation targets are surrounded by a cup-shaped double-membrane structure called a phagophore. The phagophore expands and seals to form a double-membrane autophagosome vesicle that delivers the cargo to the vacuole for degradation (Soto-Burgos et al. 2018). Autophagy is mediated by a group of core *AUTOPHAGY-RELATED* (*ATG*) genes, and disrupting core *ATG* genes usually causes the disruption of the entire autophagic process (Li and Vierstra 2012; Yang et al. 2021). In plants, this has been most studied in Arabidopsis, but has now also been shown to be conserved in other plant species, including crops (Tang and Bassham 2018; Agbemafle et al. 2025; Gross et al. 2025).

Autophagy is active at a basal level under normal conditions, is highly upregulated upon environmental stress, and can be induced by developmental stimuli. Autophagy affects many stages of plant development, such as vegetative growth (Wada et al. 2015; Minina et al. 2018; Feng et al. 2022; Goh et al. 2022), vascular tissue development (Kwon et al. 2010; Escamez et al. 2016), flowering (Kurusu et al. 2014; Hu et al. 2022), pollen development (Kurusu et al. 2014; Lu et al. 2025) and fruit ripening (Sánchez-Sevilla et al. 2021; Kumaran et al. 2025). In maize, autophagy has been shown to play a role in responses to nitrogen deficiency, fixed carbon starvation and thermotolerance (McLoughlin et al. 2018, 2020; Ma et al. 2025; Tang et al. 2025).

Autophagy is also crucial during seed development, with seed yield dependent on the level of autophagy (Guiboileau et al. 2012; Minina et al. 2018). Autophagy functions in two potential pathways during seed development, that of nutrient remobilization in maternal leaves, and in the developing seed itself. In both Arabidopsis and maize, impaired autophagy causes altered nitrogen mobilization from the mother plant to the seed, resulting in changes in seed C/N ratio (Guiboileau et al. 2012; Li et al. 2015). Autophagy is active in developing Arabidopsis seeds, and Arabidopsis *atg* mutants have altered storage protein content and processing (Di Berardino et al. 2018). In addition, *atg* mutant seeds of Arabidopsis are smaller in size than those of WT (Barros et al. 2017; Minina et al. 2018). Reciprocal crosses demonstrated that the maternal genotype affects seed protein content and germination, but not seed size (Erlichman et al. 2023), and tissue-specific complementation of an *atg5* autophagy mutant demonstrated that autophagy active only in the seed can rescue defects in seed composition but not seed size and yield (Marmagne et al. 2024). In contrast, autophagy is active in developing maize endosperm (Arcalís et al. 2022), but despite having reduced yield (Li et al. 2015), maize *atg* mutants have normal starch and storage protein content in developing endosperm (Barros et al. 2023). Instead, these mutants have an altered metabolome related to oxidative stress and accumulation of damaged mitochondria, suggesting that autophagy is important in protecting cells against damage during seed development.

These observations raise critical questions of the extent of and reasons for the differences between the role autophagy plays in seeds in maize compared with the better studied Arabidopsis, and the role of maternal versus seed autophagy in maize yield. To address these questions, we here evaluated the effect of loss of autophagy in *atg10* mutants on maize reproductive development. We found that flowering is significantly delayed in *atg10* mutants and that these mutants have reduced yield due to both smaller size and decreased number of kernels. These effects were almost entirely dependent on the maternal genotype rather than that of the embryo, suggesting that they may be controlled by nutrient remobilization from maternal tissue rather than autophagy in the embryo or endosperm.

## Results

### Flowering is delayed in *atg10* mutants

We have previously identified two independent maize mutants with decreased expression of *ATG10*, from the UniformMu (McCarty et al. 2005) and Ac/Ds collections (Vollbrecht et al. 2010), and also generated a *trans*-heterozygote of these mutant alleles. These mutants have minor deficiencies in vegetative growth and defects in autophagy and stress responses (Tang et al. 2025). Autophagy can affect flowering time by degrading flowering regulatory proteins (Hu et al. 2022), and overexpressing Arabidopsis *ATG8* or apple *ATG7b* accelerated flowering (Slavikova et al. 2008; Wang et al. 2017). In rice, *atg5* and *atg7* mutants showed delayed heading and flowering (Kurusu et al. 2014; Hu et al. 2022). To test whether flowering time is affected by autophagy in maize, days to anthesis was measured within the segregating populations of *atg10-Mu*, *atg10-Ds* and *atg10-MuDs*. In *atg10-Mu* and *atg10-MuDs* populations, plants with two mutant alleles flowered 4.5 d and 1.5 d later than siblings with two WT alleles, respectively (Figure S1). These results suggest that autophagy plays a minor role in determining flowering time in maize.

**Figure S1.**
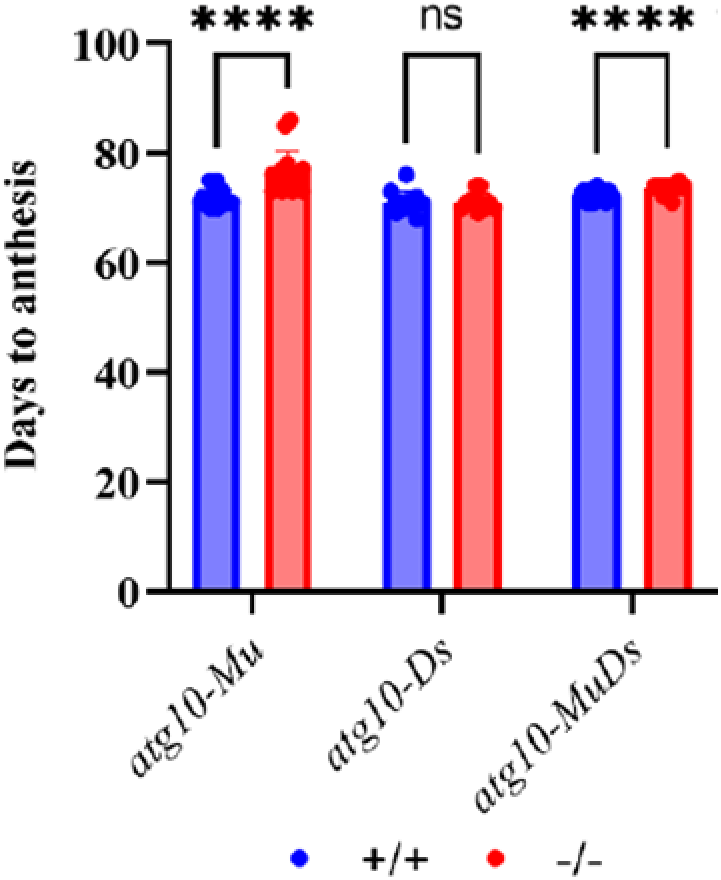
*atg10* mutants have delayed flowering. Quantification of days to anthesis for the indicated genotypes. Values are means ± SD (10<n<15). Dots in the figure indicate individual data points. Siblings with two wild-type alleles and siblings with two mutant alleles from segregating populations are compared. Asterisks indicate significant differences determined by Student’s t test (**p<0.01, ****p<0.0001, ns-not significant).

### Grain yield is reduced in *atg10* mutants

Autophagy is a major degradation and recycling pathway, and plays an important role in remobilizing nutrients from senescing tissues to developing tissues including seeds (Li et al. 2015). Reduced seed yield has been shown for autophagy mutants in various plant species including Arabidopsis (Minina et al. 2018; Yu et al. 2019; Zhen et al. 2019, 201, 2021; Fan et al. 2020; James et al. 2025) and maize (Li et al. 2015).

Kernel size and weight are two important components of maize yield (Liu et al. 2017), and we hypothesized that these parameters may be reduced in *atg10* mutants. Kernels produced by plants with two mutant alleles had visibly reduced width and length in all three mutant allele combinations when compared to kernels produced by their WT siblings (Figure 1A). Kernel length and kernel width were significantly reduced by up to 20% in *atg10* mutant lines relative to WT (Figure 1B and 1C). 100-kernel weight was also significantly reduced in kernels produced by *atg10* plants (Figure 1D).

**Figure 1.**
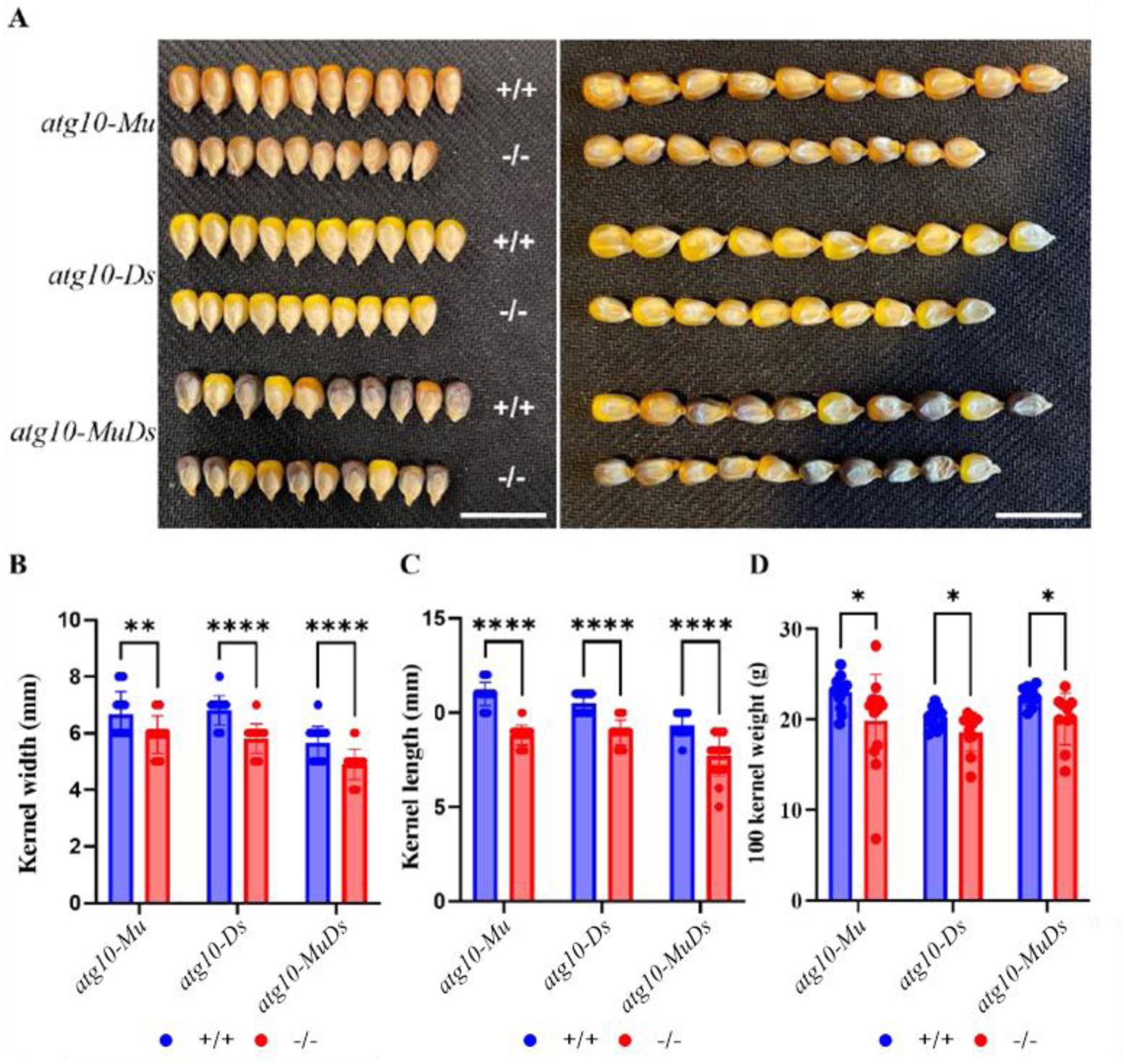
Kernel size is reduced in *atg10* mutants. **A.** Kernel phenotypes of *atg10-Mu*, *atg10-Ds* and *atg10-MuDs*. Kernels harvested from plants with two wild-type alleles (+/+) and plants with two mutant alleles (-/-) are shown. Scale bar = 1 cm. **B-D.** Quantification of kernel width (n=30) (B), kernel length (n=30) (C) and 100-kernel weight (10<n<15) (D). Values are means±SD. Dots in the figure indicate individual data points. Siblings with two wild-type alleles and siblings with two mutant alleles from segregating populations are compared. Asterisks indicate significant differences determined by Student’s t test (*p<0.05, **p<0.01, ****p<0.0001).

Ear length is another important component of maize grain yield and increased ear length leads to increased kernel number, which finally leads to increased yield (Huo et al. 2016; Jia et al. 2020; Ning et al. 2021; Luo et al. 2022). While no obvious difference in grain-fill was observed when comparing ears harvested from WT plants and ears harvested from *atg10* plants (Figure 2A), in mature ears produced by *atg10* plants, ear length and kernel number per row were all markedly reduced relative to ears harvested from WT plants (Figures 2A-2C), leading to reduction in total kernel number per plant by 13.5%-32.5% (Figure 2D). *ATG10* mutation resulted in a substantial reduction in ear weight (Figure 2E), and reduction in total kernel weight per plant by up to 37% (Figure 2F). Altogether, these results indicate that autophagy is required for kernel size and yield.

**Figure 2.**
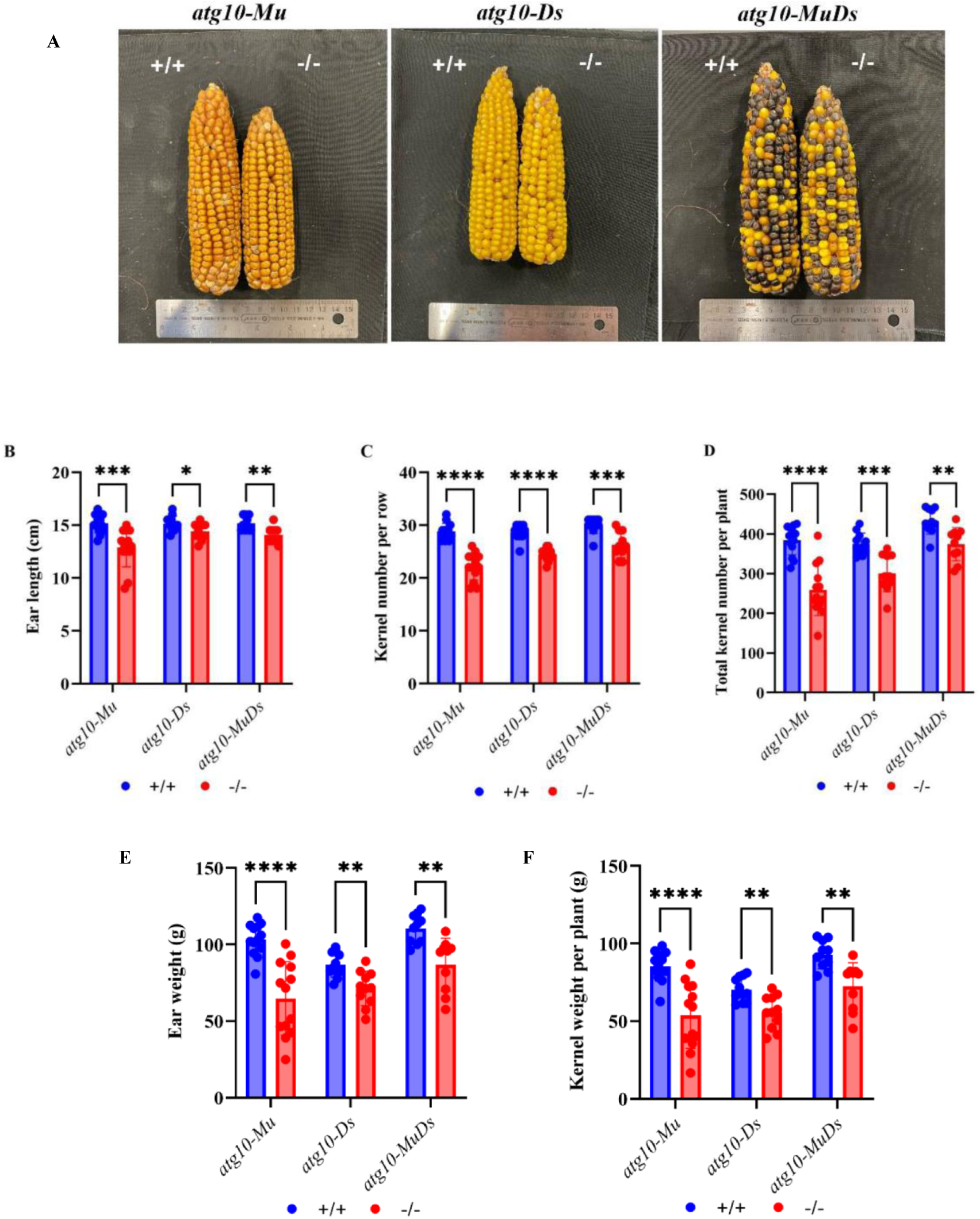
Ear length and kernel number are reduced in *atg10* mutants. **A.** Representative images comparing ears produced by a plant with two wild-type alleles with ears produced by a plant with two mutant alleles. **B-F.** Quantification of ear length (B), kernel number per row (C), total kernel number per plant (D), ear weight (E), and kernel weight per plant (F). Values are means±SD (10<n<15). Dots indicate individual data points. Ears harvested from siblings with two wild-type alleles and siblings with two mutant alleles from segregating populations are compared. Asterisks indicate significant differences determined by Student’s t test (*p<0.05, **p<0.01, ***p<0.001, ****p<0.0001).

In maize, endosperm is a triploid tissue that occupies most of the mature grain and provides nutrients during seed germination and seedling growth (Li and Berger 2012; Dai et al. 2021). Transcripts of many *ATG* genes are upregulated in maize endosperm during seed development (Li et al. 2015), and an increase in both ATG8 and ATG8-PE was observed in endosperm starting at 14 DAP (Chung et al. 2009). We hypothesized that the reduction in kernel size and kernel weight in *atg10* mutants might be due to defects in endosperm development. However, light microscopy analysis of mature *atg10* endosperms revealed no structural defects in the aleurone or starchy endosperm and showed a normal packing of starch granules (Figure S2). Zein protein content and non-zein protein content were analyzed by SDS-PAGE and no obvious differences were observed (Figure S3). It is therefore unlikely that the changes in kernel size in the *atg10* mutants are due to gross developmental defects.

**Figure S2.**
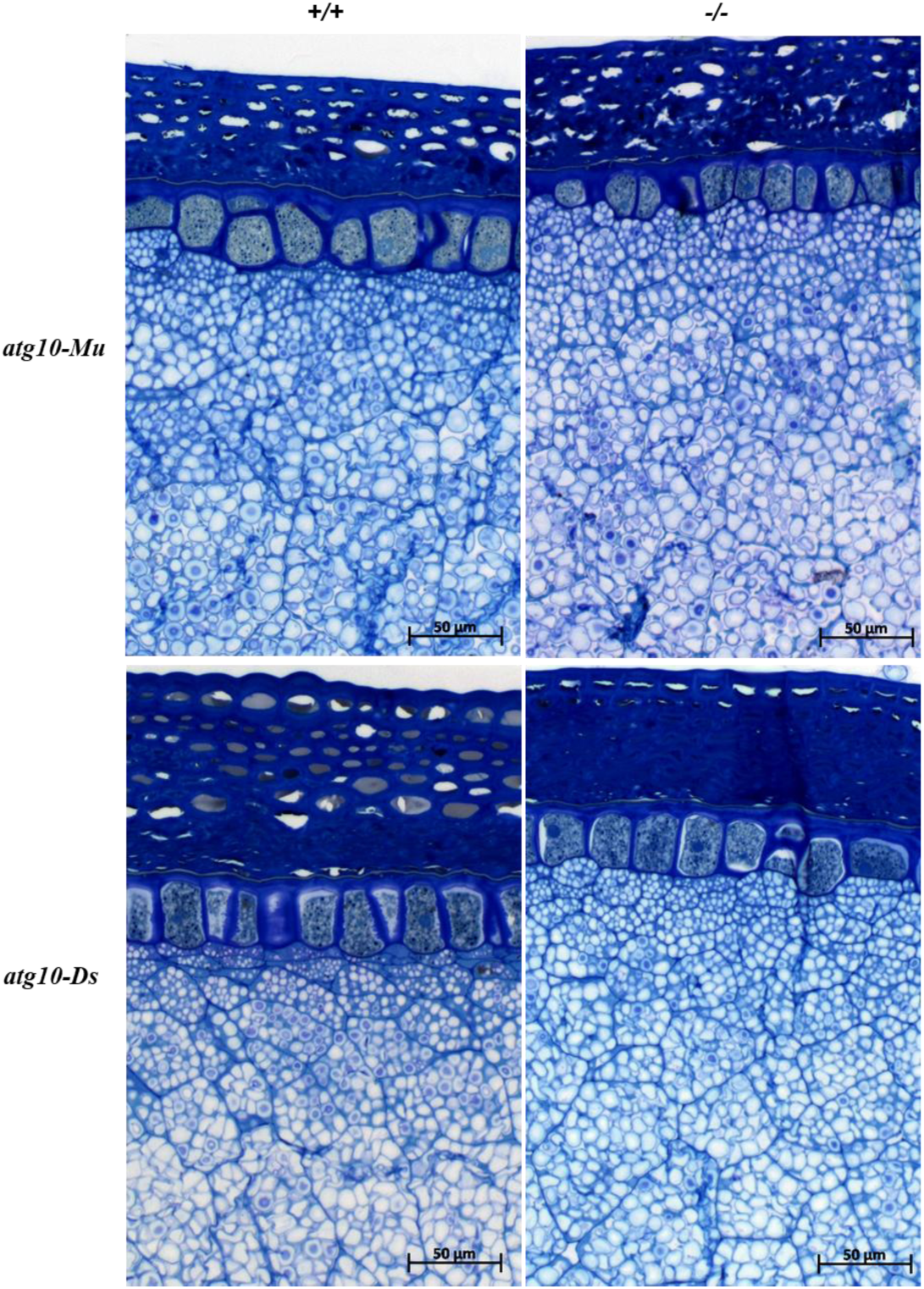
Light microscopy analysis of mature endosperms shows normal mutant kernel structure. Mature kernels were fixed, embedded, sectioned, stained with Toluidine blue and basic fuschsin, and visualized by light microscopy. Scale bars = 50 µm

**Figure S3.**
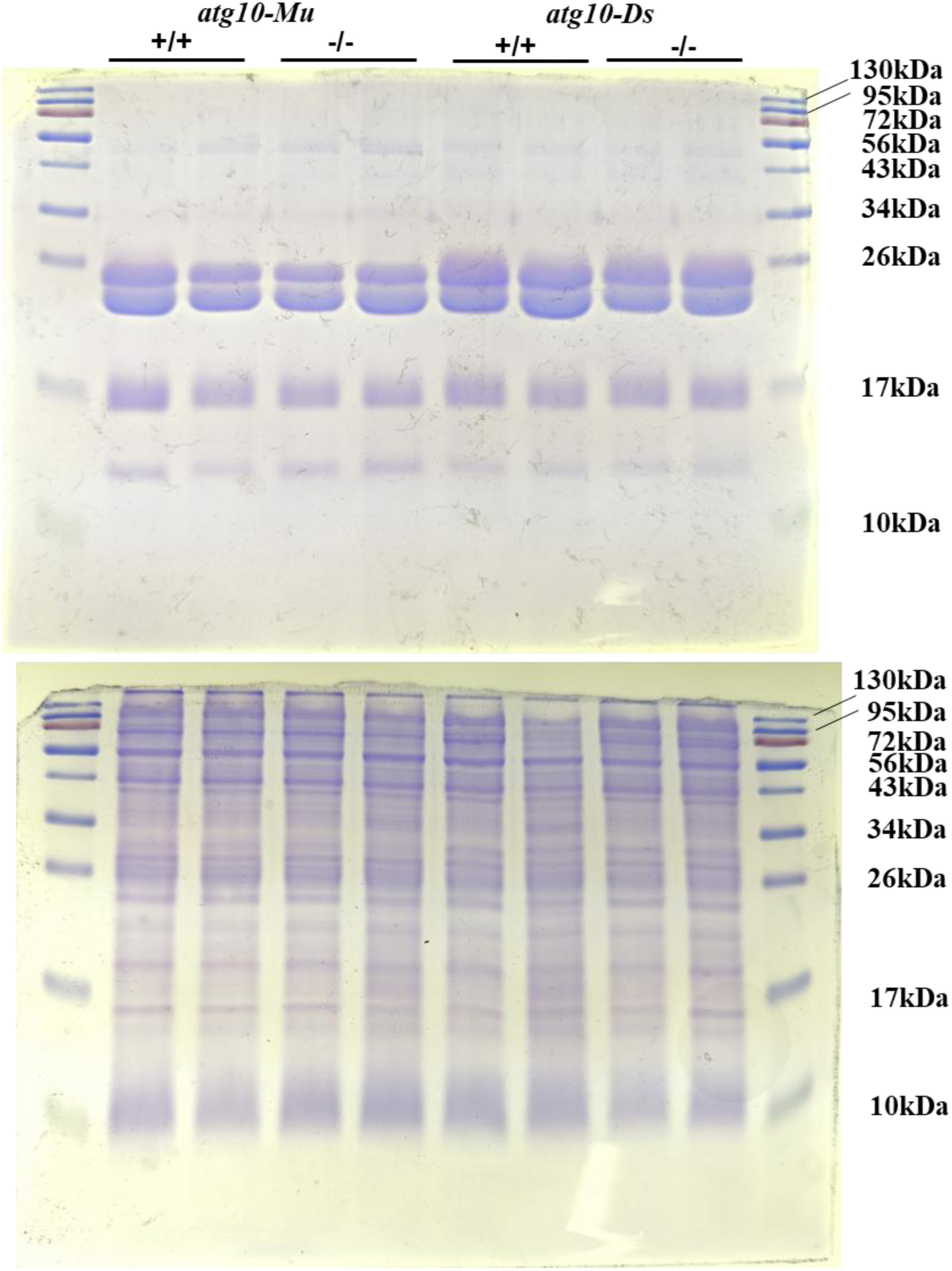
SDS-PAGE analysis of zein and non-zein proteins. Zein (upper panel) and non-zein (lower panel) proteins were extracted from 50 mg kernel powder. Protein samples were separated by SDS-PAGE and stained with Coomassie blue. The relative molecular mass of the protein ladder is indicated.

### Maternal plants contribute to the ear and kernel phenotypes in *atg10* mutants

We considered two non-exclusive hypotheses: (1) The kernel phenotype seen in the *atg10* mutants is due to defective autophagy in the kernels themselves, either endosperm and/or embryo (Barros et al. 2023; Erlichman et al. 2023; Marmagne et al. 2024) or (2) the kernel phenotype is due to loss of autophagy in the mother plant, due to defects in nutrient remobilization (Wada et al. 2009; Guiboileau et al. 2012; Li et al. 2015).

To test these hypotheses, we first self-pollinated heterozygous plants and compared the weight and size between mutant and WT kernels developing on the same ear (Figure 3A). However, no difference was observed in kernel weight, length or width between any mutant allele combination and the WT (Figure 3B-D), indicating that the seed genotype, either endosperm or embryo, does not affect these phenotypes.

**Figure 3.**
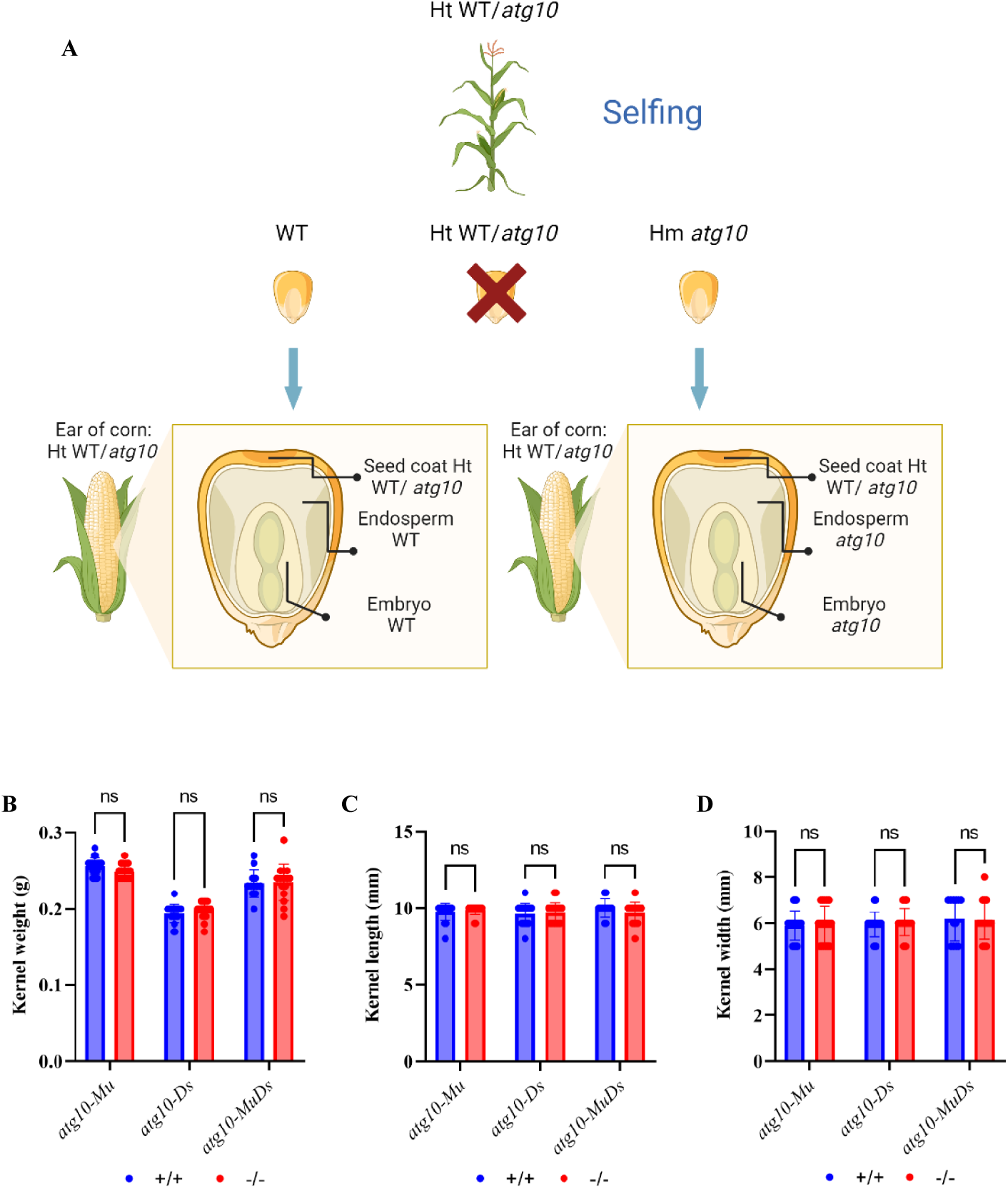
Segregating kernels on the same ear have the same phenotype. **A.** Schematic of crossing strategy to generate kernels with different genotypes developing on the same ear. **B-D.** Quantification of kernel weight (B), kernel length (C) and kernel width (D). Values are means±SD (10<n<20). Dots indicate individual data points. Statistical analysis was performed using Student’s t test (ns-not significant).

We next hypothesized that maternal plants may play a role in determining the kernel phenotype of *atg10* mutants. To address this hypothesis, reciprocal crosses were conducted between WT siblings and *atg10-Mu* siblings within the same segregating population, resulting in kernels with heterozygous embryos developing on either WT or mutant ears (Figure 4). Note that the triploid endosperm contained either one or two mutant *atg10* alleles, depending on whether the mutant allele was derived from the paternal or maternal plant. As the endosperm genotype did not affect seed phenotypes (Figure 3), an effect of the endosperm *ATG10* copy number is unlikely.

**Figure 4.**
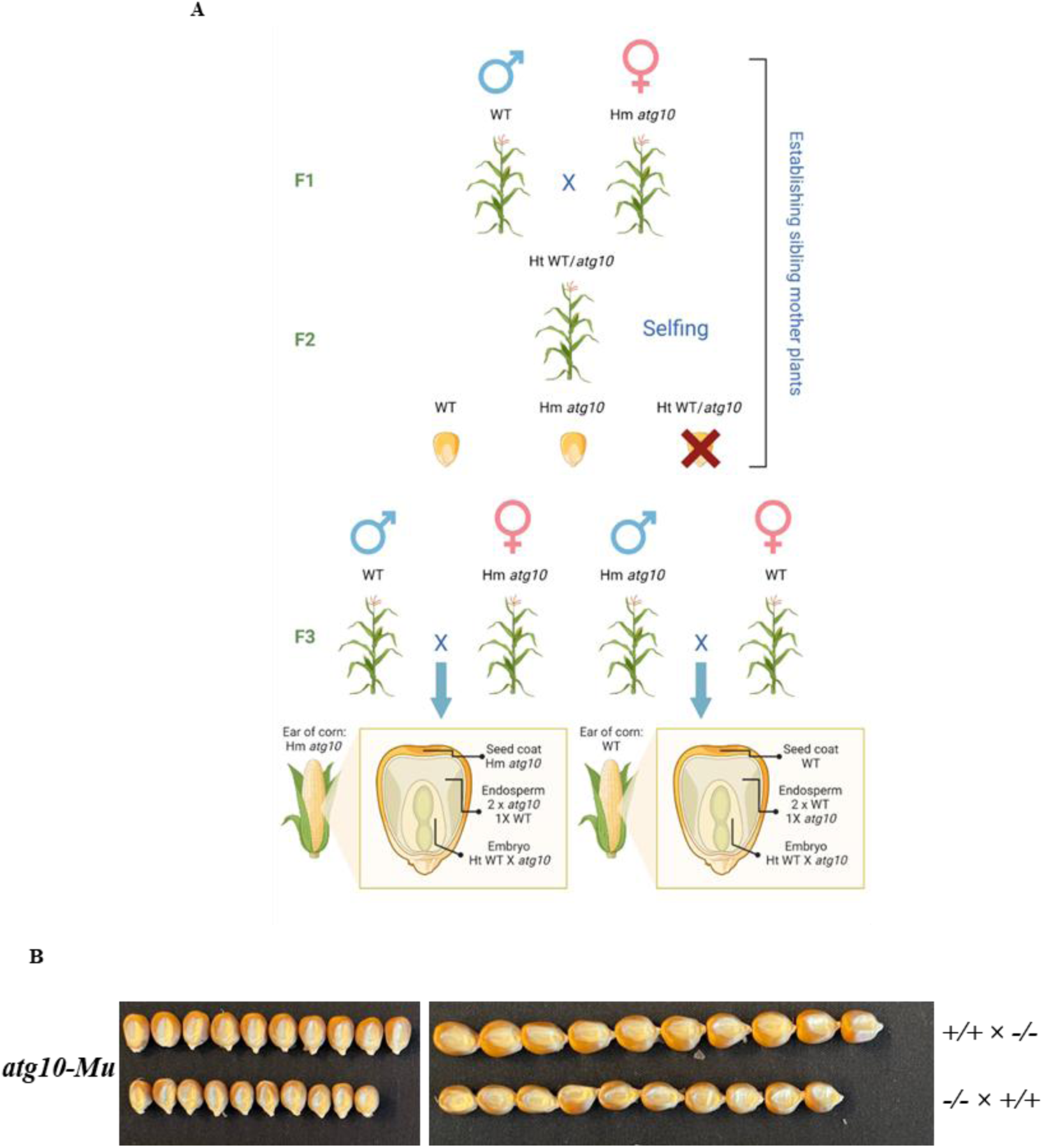
Reciprocal crosses show the contribution of the maternal plant genotype to *atg10* kernel phenotype. **A**. Schematic of reciprocal crossing strategy to generate kernels with the same genotypes developing on ears of distinct maternal genotype. **B.** Images of kernels harvested from plants with two wild-type alleles (+/+) and plants with two mutant alleles (-/-). Genotype before the cross marker indicates the maternal genotype. Scale bar = 1cm.

A significant influence of the maternal genotype on kernel size and kernel weight was observed (Figure 5). Kernels produced by a WT mother were larger and heavier than kernels produced by a mutant mother (Figure 5A-C). In addition, ear length, kernel number per row, and total kernel number per plant were significantly elevated in WT mother relative to mutant mother plants (Figure 5D-F). The differences in kernel size, weight and number also caused a significant difference in yield, measured by ear weight and total kernel weight per plant (Figure 5G, H). Taken together, these results suggest that the maternal plant genotype, rather than the embryonic genotype, is the major contributor to the kernel, ear and yield phenotype.

**Figure 5.**
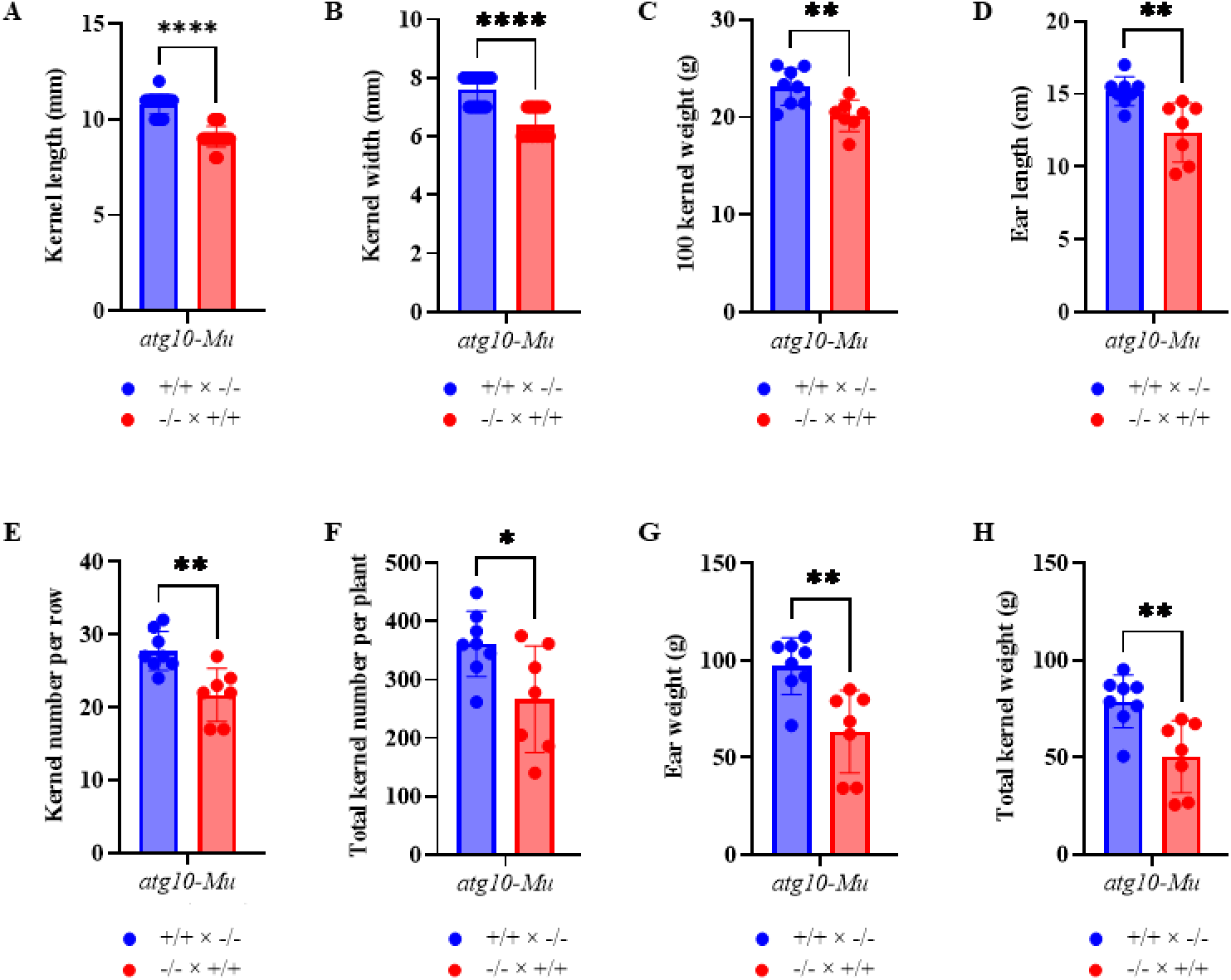
Maternal plant contributes to the kernel, ear and yield phenotypes in *atg10*. **A-H.** Quantification of kernel length (n=20) (A), kernel width (n=20) (B), 100-kernel weight (n=8) (C), ear length (n=8) (D), kernel number per row (n=8) (E), total kernel number per plant (n=8) (F), ear weight (n=8) (G) and total kernel weight per plant (n=8) (H). Values are means±SD. Dots in the figure indicate individual data points. Asterisks indicate significant differences determined by Student’s t test (*p<0.05, **p<0.01, ****p<0.0001).

## Discussion

Autophagy functions at multiple stages of a plant’s life cycle, both in developmental processes and in responses to changes in environmental conditions. We show here that autophagy plays a minor role in determining flowering time in maize, as flowering is slightly delayed in *atg10* mutants. This phenotype is consistent with effects on flowering observed in other plant species (Slavikova et al. 2008; Kurusu et al. 2014; Wang et al. 2017; Hu et al. 2022). Autophagy also plays a significant and substantial role in grain yield. Kernel weight, kernel size, ear size and kernel number were all reduced in *atg10* mutants, leading to a reduction in total grain yield. Reciprocal crosses indicated that the maternal plant genotype is the major contributor to the kernel and ear phenotypes in *atg10* mutants, rather than the genotype of the embryo, suggesting an important role for nutrient mobilization from maternal tissue to seeds.

Autophagy is known to be an important factor in nitrogen remobilization from senescing tissues to seeds, and plant autophagy mutants have reduced nitrogen remobilization from leaves to seeds as well as reduced nitrogen use efficiency (Guiboileau et al. 2012; Li et al. 2015; Yu et al. 2019; Zhen et al. 2019, 2021; Fan et al. 2020). This could explain the reduced kernel size, kernel weight, and yield in *atg10* mutants. We did not see any obvious changes in protein content and composition in *atg10* kernels. The delivery of storage proteins from the endoplasmic reticulum to protein storage vacuoles in maize aleurone cells is mediated by microautophagy rather than macroautophagy, and is independent of the ATG8 lipidation pathway (Ding et al. 2022). This protein transport pathway may therefore be functional in *atg10* mutants (Phillips et al. 2008). Another factor contributing to the yield reduction in *atg10* mutants is a reduction in ear length and kernel number. Kernel number is primarily affected by the growth and biomass accumulation of the ear, and ear biomass is determined by the maternal plant biomass accumulation, as well as the allocation of biomass from the plant to the ear (Borrás and Vitantonio-Mazzini 2018). Autophagy is known to remobilize nutrients from older tissues to younger tissues and support biomass production (Wada et al. 2015). The reduction in ear length and kernel number in *atg10* mutants may therefore also be due to a deficiency in nutrient remobilization. Interestingly, the kernel number of maize *atg12* mutants grown in the greenhouse, or in fields pre-treated with fertilizer, was not significantly different from WT grown under the same condition (Li et al. 2015). However, under nitrogen deficiency, *atg12* ear development was significantly stunted relative to WT (Li et al. 2015). Thus, it is possible that environmental stress such as heat or nutrient deficiency contributes to the reduction in ear length and kernel number in field-grown *atg10* mutants.

We considered that the reduced yield of *atg10* mutants could be caused by the loss of autophagy in maternal tissues, in the embryo, the endosperm, or both. To address this, we first generated plants in which the maternal plant was heterozygous for the mutant allele, and the kernel genotypes were segregating within the same ear, as illustrated in Figure 3A. A comparison of mutant and WT kernels derived from the same ear indicated no difference in size or weight, and the kernel phenotype therefore appears independent of its genotype. We next made reciprocal crosses, such that the maternal plant was either WT or homozygous for the mutant allele, and the seeds were all heterozygous, shown in Figure 4A. In this situation, kernels derived from a mutant maternal plant were smaller than those from a WT, suggesting that the maternal autophagy status controls the size of the seeds.

While it is known that autophagy mutants in some plant species tend to have smaller and fewer seeds than the corresponding WT (Guiboileau et al. 2012; Li et al. 2015; Barros et al. 2017; Minina et al. 2018), the strong effect of maternal genotype on seed size and yield was unexpected, and distinct from effects seen in Arabidopsis. In Arabidopsis, maternal genotype primarily affects seed protein content and germination, rather than seed size, which is determined by autophagy activity within the developing seed. (Erlichman et al. 2023; Marmagne et al. 2024). These differences were unexpected, and one possibility is that they are due to differences in seed development processes in maize when compared with Arabidopsis (Gillmor et al. 2020). In Arabidopsis the endosperm is largely consumed during seed development, with the mature seed mainly consisting of the embryo with a single aleurone cell layer. By contrast, in maize the endosperm is maintained in the mature seed, and the bulk of the seed consists of the starchy endosperm. Mechanisms determining seed size are therefore likely to differ between the two species.

In conclusion, maize *atg10* mutants showed delayed flowering and reduced grain yield, due to a reduction in kernel size, weight and number. The maternal plant genotype is the major factor contributing to these kernel phenotypes, and we hypothesize that this is due to decreased nutrient remobilization from maternal tissue to developing seeds. Maize seed size has been suggested to be controlled maternally, although the mechanistic basis is likely to be complex (Zhang et al. 2016). Our results suggest that autophagy may be one factor contributing to this maternal control.

## Materials and methods

### Plant materials and growth conditions

Autophagy mutant identification and characterization were previously described in (Tang et al. 2025). Insertion allele *atg10-Mu* (mu1052008) was derived from the UniformMu collection (McCarty et al. 2005) and was obtained from the Maize Stock Center (http://maizecoop.cropsci.uiuc.edu). Insertion allele *atg10-Ds* (I.S06.1765) was generated by the Ac/Ds project (Vollbrecht et al. 2010). Both lines were backcrossed to W22 at least three times. BC_3_F_2_ segregating populations were used in this study. A trans-heterozygous *atg10-Mu*/*atg10-Ds* allele (named *atg10-MuDs*) was generated by crossing *atg10-Mu/+* with *atg10-Ds/+*.

Plants were grown at the Iowa State University Curtiss Research Farm under well-watered conditions. Three different BC_3_F_2_ segregating populations per mutant allele were planted in three different four-row blocks, respectively, following a randomized block design. 60 plants were planted in each block. The distance between two rows was 60 cm and the distance between two plants in the same row was 20 cm. Leaf samples were taken for genomic DNA extraction and genotyping.

### Measurements of plant traits

Flowering time was measured as the number of days from planting to the day when plants were shedding pollen. For ear and kernel traits, harvested ears were first air dried to constant weight before measurements. Traits measured included ear weight, total kernel weight per plant, 100-kernel weight, kernel length, kernel width, ear length, number of kernels per row, kernel row number per ear and total kernel number per plant. Ear weight is the weight of whole dry ears with kernels attached.

For traits of segregating kernels from the same ear, individual kernels were first used for measurements of weight, length, width, and then germinated to seedlings. Leaf tissue was collected for genomic DNA extraction and genotyping.

### Genotyping

Genomic DNA extraction was done as described previously (Dietrich et al. 2002). Leaf tissue was ground in CTAB extraction buffer (0.1 M Tris, 0.7 M NaCl, 10 mM EDTA, 1% CTAB (w/v), 1% 2-mercaptoethanol (v/v)). The lysate was incubated at 65°C for 30 min with mixing by inverting the tube every 10 min. After cooling down to room temperature, the lysate was mixed with 400 µL chloroform/iso-amyl alcohol (24:1) by vortexing for 10∼15 seconds. The mix was centrifugated at room temperature for 10 min at 10,000 *g*. The supernatant was transferred to a new tube and mixed with 2.5 volumes 100% ethanol, then incubated at −20°C for at least one hour (or overnight). After incubation, the tube was centrifugated at room temperature for 15 min at 10,000 *g*. The supernatant was discarded, and the pellet was washed with 500 µL 70% ethanol, followed by a centrifugation at room temperature for 5 min at 10,000 *g*. This step was repeated twice. After the final wash, the tube was air dried, and 50 µL ddH_2_O was added to dissolve the DNA. Genotyping PCR was done using GoTaq G2 Flexi (Promega, M7801) as previously described (McCarty et al. 2013). Primers used are listed in Table S1.

### Zein and non-zein protein analysis

Zein and non-zein protein measurement was performed as previously described (Wu and Messing 2012; Lappe et al. 2018). Kernels were wrapped in four layers of aluminum foil and pulverized with a hammer, then ground into powder with a mortar and pestle. 50 mg of flour was transferred to a 2 mL tube and mixed with 400 µL zein extraction buffer (70% ethanol/2% 2-mercaptoethanol (v/v)) by vortex. The mix was kept at room temperature overnight and then centrifugated at 10,000 *g* for 10 min at room temperature. The supernatant was used for zein extraction, and the pellet was used for non-zein extraction.

For zein extraction, 100 µL supernatant was transferred to a new tube and mixed with 10 µL 10% (w/v) SDS. The mixture was dried by vacuum and resuspended in 100 µL of distilled water.

For non-zein extraction, the pellet was resuspended in 400 µL zein extraction buffer to remove zein proteins. The mix was centrifuged at 10,000 *g* for 10 min at room temperature. These steps were repeated three times. The pellet was resuspended in 400 µL non-zein extraction buffer (12.5 mM sodium borate, 5% (w/v) SDS and 2% 2-mercaptoethanol (v/v)) and kept at 37°C for two hours with occasional vortex. After incubation, the mix was centrifuged at 10,000 *g* for 10 min, and then 100 µL of the non-zein supernatant was transferred to a new tube.

For SDS-PAGE analysis, 4 µL of each sample was loaded onto a 15% SDS-PAGE gel followed by staining with Coomassie Blue.

### Microscopy

Sample preparation and microscopy was performed at the Iowa State University Roy J. Carver High Resolution Microscopy Facility. Mature kernels imbibed in H_2_O for 2 days at 50°C were immersed in fixative (2% paraformaldehyde, 3% glutaraldehyde, 0.1 M cacodylate, pH 7.2), dissected by hand with a fresh razor blade under fixative into small pieces, and then fixed for 48 hours at 4°C. Samples were washed in the cacodylate buffer for 10 mins for 3 times, and post-fixed with 1% osmium tetroxide in 0.1 M sodium cacodylate buffer for 1 hour at room temperature. Samples were washed with deionized water for 15 mins for 3 times, and en bloc stained using 2% uranyl acetate in distilled water for 1 hour. Samples were washed in distilled water for 10 mins and dehydrated through a graded ethanol series (25, 50, 70, 85, 95, 100%) for 1 hour each step. Samples were further dehydrated with 3 changes of pure acetone, 15 minutes each, and infiltrated with Spurr’s formula (hard) epoxy resin (Electron Microscopy Sciences, Hatfield PA) with graded ratios of resin to acetone until fully infiltrated with pure epoxy resin (3:1, 1:1, 1:3, pure) for 6-12 hours per step. Tissues were placed into beem capsules and were polymerized at 70°C for 48 hours. Thick sections (1.5 µm) were made using a Leica UC6 ultramicrotome (Leica Microsystems, Buffalo Grove, IL) and stained with EMS Epoxy stain (a blend of toluidine blue-O and basic fuchsin).

### Statistical analysis

All statistical tests were done with GraphPad Prism 9. Student’s t-test was used for comparison between two independent groups.

## Acknowledgements

We thank Dr. Steve Howell for valuable discussions throughout the project, Tracey Stewart for assistance with microscopy, and Vishadinie Jayasinghe for help with genotyping. This work was supported by grants from the Binational Agricultural Research and Development Fund (IS-5760-25 to TAW and DCB) and from the National Science Foundation (Plant Genome Research Program IOS 1444339 to DCB). Schematic diagrams were created with BioRender.com.

---

**Supplemental table 1.**
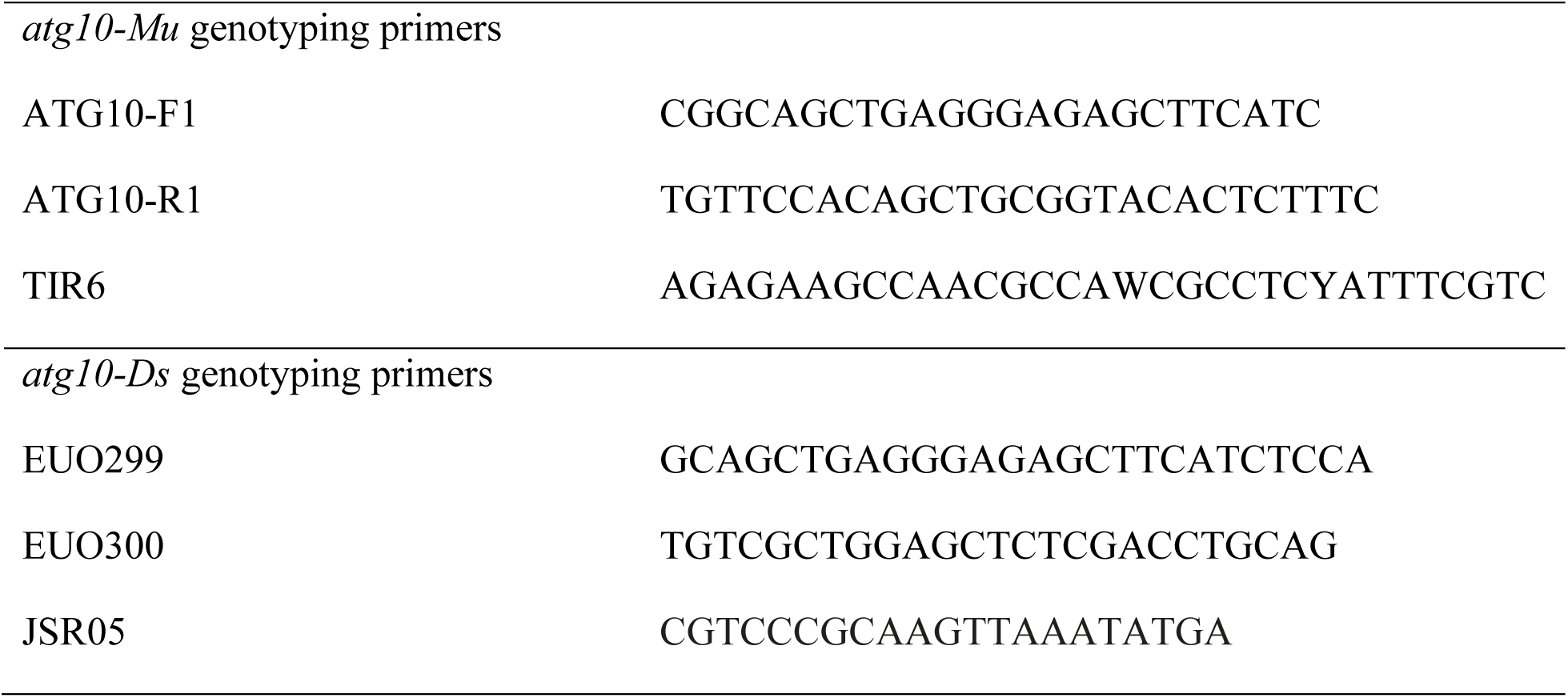
Primers used in this study.

